## Supplemental Data for "A Chaperonin Complex Regulates Organelle Proteostasis in Malaria Parasites"

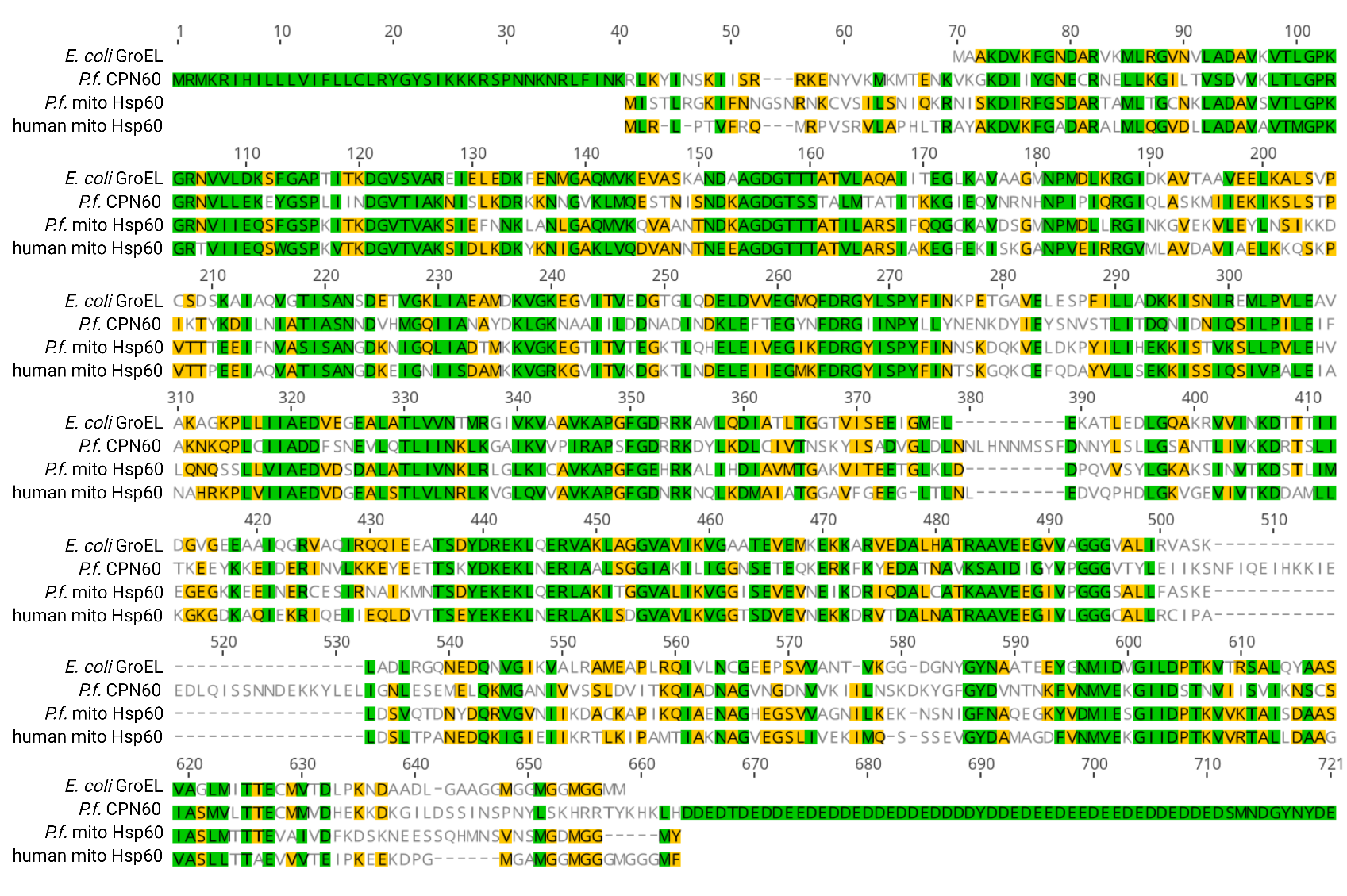
S1 Fig

**S1 Fig.** Amino Acid sequence alignment between (1) *E. coli* GroEL (UNIPROT P0A6F5), (2) *Plasmodium falciparum* apicoplast CPN60 (PF3D7_1232100), (3) *Plasmodium falciparum* mitochondrial Hsp60 (PF3D7_1015600), and (4) human mitochondrial Hsp60 (UNIPROT P10809). Green represents Identity and high similarity (>80%) and Yellow similarity (>60%). Note the N-terminal extension of CPN60 representing the transit peptide which is removed upon apicoplast localization. Additional extensions appearing only in CPN60 are two inner loops and the long C-terminal stretch, which may interfere with PBZ binding.

S2 Fig

**A.**

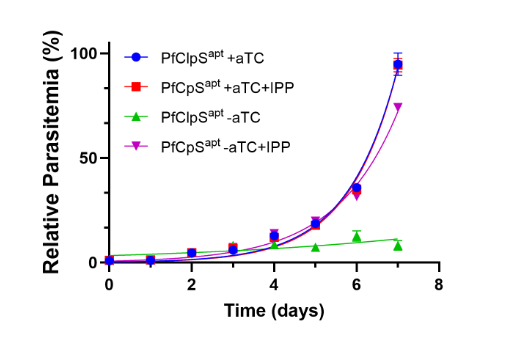

| **aTC conc.** | **500 nM** | | **16 nM** | | **8 nM** | | **4 nM** | |
| --- | --- | --- | --- | --- | --- | --- | --- | --- |
| temperature | 37^o^c | 40^o^c | 37^o^c | 40^o^c | 37^o^c | 40^o^c | 37^o^c | 40^o^c |
| Doubling Time | 1.374 | 1.581 | 1.473 | 1.785 | 1.876 | 1.964 | 2.955 | 3.496 |

**B.**

**S2 Fig. A.** PfClpS^apt^ parasites were cultured in the presence or absence of aTC and IPP, and parasitemia was measured every 24 hours for 7 days using flow cytometry. Parasites lacking aTC display a growth defect beginning on day 4, followed by a decline in parasitemia. The addition of IPP rescues the growth defect, indicating apicoplast dysfunction. Parasitemia was normalized to the maximum value observed under aTC treatment on day 7, and Normalized data are represented as mean ± SEM for three technical replicates. **B.** CPN60^V5-apt^ parasites were washed and incubated with different aTC concentrations (8 nM, 4 nM, 16 nM and 500 nM), subjected to HS and then allowed to grow at 37^o^C for three days, while being measured daily by flow cytometry. Data were fit to an exponential (Malthusian) growth curve (graph shown on Fig. 4F) and the doubling time as calculated is shown here.

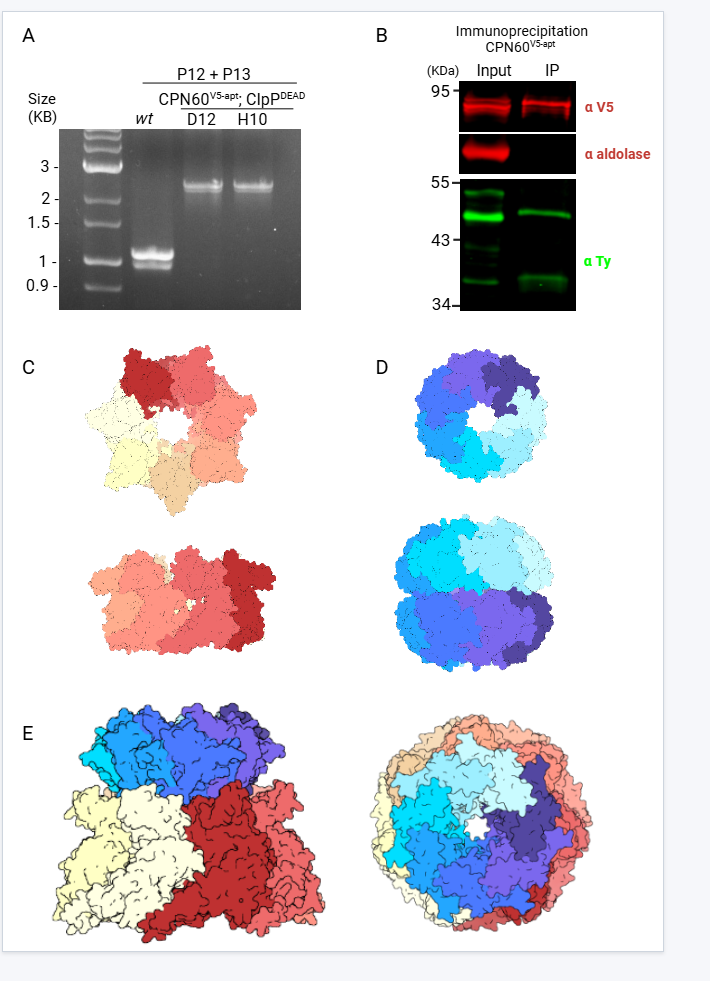
S3 Fig.

**S3 Fig. A.** Genotyping by PCR to confirm ClpP^DEAD-Ty^ integration at the *hsp110* locus. Genomic DNA was purified from transfected isolated parasites clones (D12 and H10), and primers P12 and P13 were used to specifically amplify the integrated region. A shift of 1000 bp corresponds to the integration of ClpP^DEAD-Ty^. **B.** Co-IP of CPN60^V5-apt^. Parasites were isolated and sonicated, and extracts were incubated with anti-V5 antibody-conjugated beads (for CPN60 pulldown) Input and IP samples were loaded on SDS-page and blotted with anti-Ty, anti-V5 antibodies and anti-aldolase as a negative control. **C.** The X-ray structure of apicoplast CPN60 (PDB ID: 7K3Z)^31^. The different chains are highlighted in different yellow and red colors, revealing the heptameric arrangement of the CPN60 ring. On the left is a top view, and on the right is a side view. **D.** The X-ray structure of PfClpP (PDB ID: 2F6I)^22^. The different chains are highlighted in different blue shades, revealing the heptameric arrangement of the PfClpP ring. On the left is a top view, and on the right is a side view. **E.** AF3 structure prediction of the interaction between the two heptameric rings of CPN60 and PfClpP. The PfClpP ring is shown tilting upward, binding to the upper side of CPN60. PfClpP is depicted in blue and teal shades, while CPN60 is coloured in red and beige. On the left is a side view, and on the right is a top view.

S4 Fig.

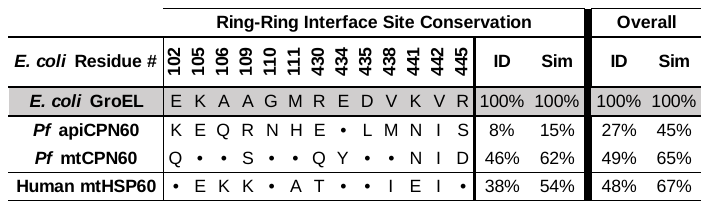

**S4 Fig.** Overall GroEL/CPN60/HSP60 and ring-ring interface binding site residue conservation. Residues that were observed to interact with ligand in the GroEL: PBZ-1587 cryoEM structure are shown in grey, with corresponding residues from sequence alignments for *P. falciparum* apiCPN60, *P. falciparum* mtCPN60, and human mtHSP60 shown below. Conserved residues are represented as dots. Percent identical (ID) and similar (Sim) residues for the ring-ring interface and overall chaperonin sequences are shown to the right.

S5 Fig.

**A.**

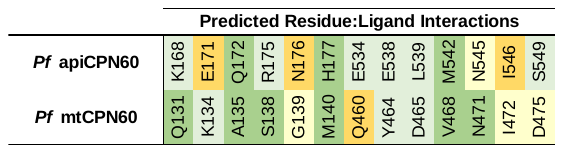

**B.**

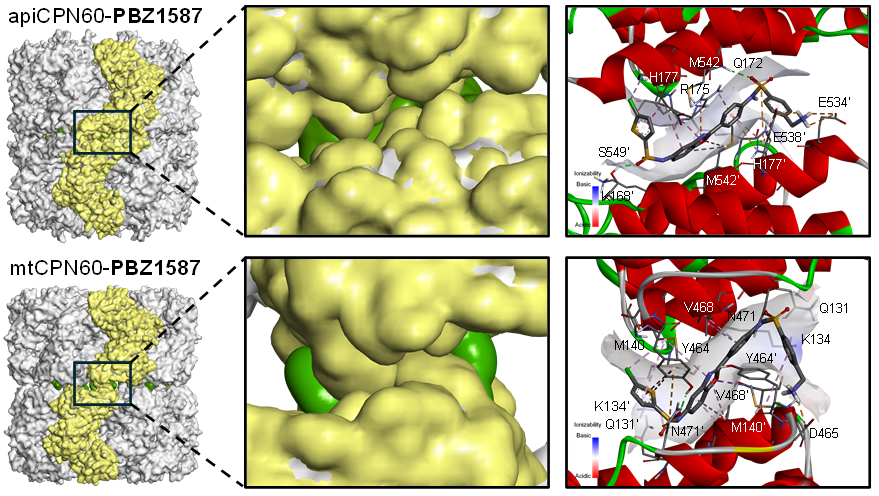

**S5 Fig.** **A.** Residue: ligand interaction map as predicted from analysis of the apiCPN60: PBZ-1587 and mtCPN60:PBZ1587 homology models. Color-coding indicates the degree of interaction: Orange = residue more distant with no appreciable interaction; Yellow = residue adjacent but no significant interaction; Light Green = positive interaction with corresponding residue from one ring; Dark Green = positive interactions with corresponding residues from both rings. **B.** Images of PBZ-1587 bound in the apiCPN60 and mtCPN60 homology models.

S6 Fig.

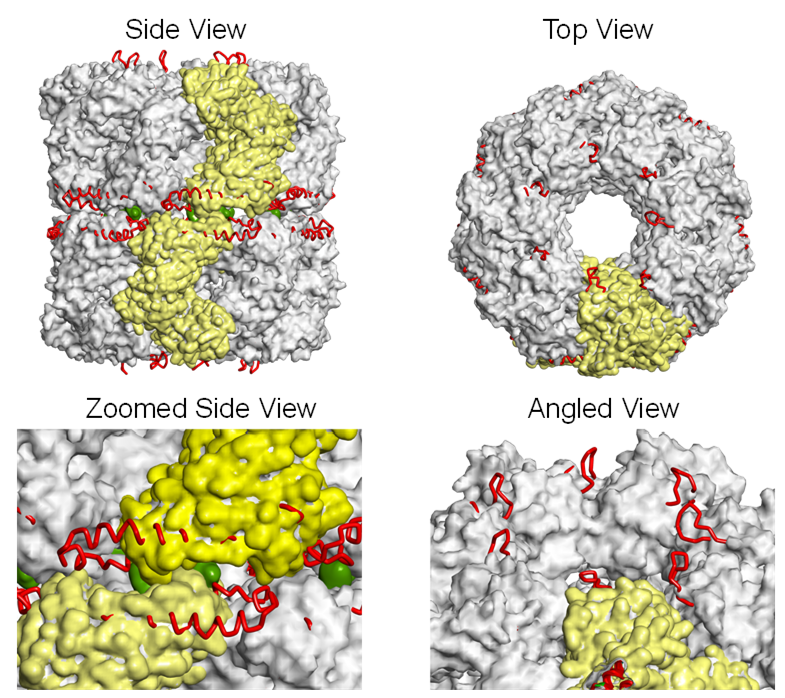

**S6 Fig.** Overlay of the apiCPN60: PBZ-1587 and mtHSP60: PBZ-1587 homology models. The mtCPN60 rings are shown as grey solvent exposed surfaces (with two individual subunits across the ring-ring interface shown in yellow), the apiCPN60 protein backbones are shown as red tubes, and bound PBZ-1587 inhibitors are shown as green surfaces. Unlike *P. falciparum* mtCPN60 and chaperonin orthologues from other species, apiCPN60 has additional loop residues extending from the apical domains (residues 370-392), and extended alpha-helices at the ring-ring interface (residues 502-529).

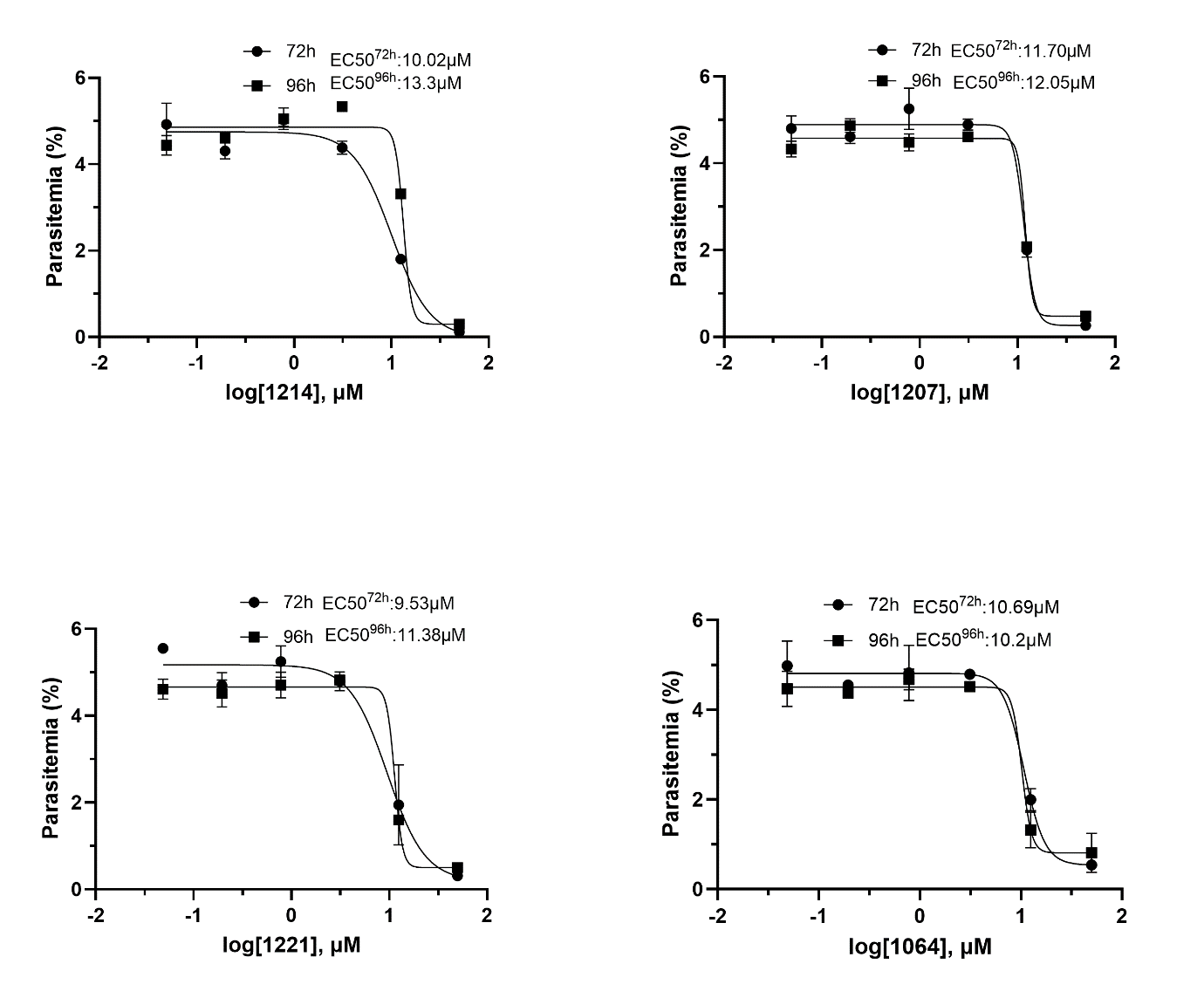
S7 Fig.

**S7 Fig. Testing phenylbenzoxazole (PBZ) GroEL inhibitors during partial CPN60 knockdown. A.** CPN60^V5-apt^ parasites were washed eight times and the incubated with 8nM aTC for 24 hours. The following days these parasites were seeded at 0.5% parasitemia and incubated in serial dilutions of PBZ compounds, ranging from 50 μM to 50 nM. Parasitemia was measured after 72 or 96 hours to calculate the drugs’ half-maximal effective concentrations (EC50). Data are fit to a dose-response equation and are represented as mean ± SEM.
